# ctDNAtools: An R package to work with sequencing data of circulating tumor DNA

**DOI:** 10.1101/2020.01.27.912790

**Authors:** Amjad Alkodsi, Leo Meriranta, Annika Pasanen, Sirpa Leppä

## Abstract

**Summary:** Sequencing of cell-free DNA (cfDNA) including circulating tumor DNA (ctDNA) in minimally-invasive liquid biopsies is rapidly maturing towards clinical utility for cancer diagnostics. However, the publicly available bioinformatics tools for the specialized analysis of ctDNA sequencing data are still scarce. Here, we present the ctDNAtools R package, which provides functionalities for testing minimal residual disease (MRD) and analyzing cfDNA fragmentation. MRD detection in ctDNAtools utilizes a Monte Carlo sampling approach to test ctDNA positivity through tracking a set of pre-detected reporter mutations in follow-up samples. Additionally, ctDNAtools includes various functionalities to study cfDNA fragment size histograms, profiles and fragment ends patterns.

**Availability:** The ctDNAtools package is freely available under MIT license at https://github.com/alkodsi/ctDNAtools.

## 1 Introduction

Molecular characterization of circulating tumor DNA (ctDNA) is finding wide applications in cancer diagnostics and clinical decision-making including, but not limited to, early cancer detection (Phallen *et al.*, 2017), tissue of origin classification (Kang *et al.*, 2017), cancer genome characterization (Parikh *et al.*, 2019), epigenetic profiling (Snyder *et al.*, 2016), and cancer monitoring after treatment (Chaudhuri *et al.*, 2017). Although analyzing cfDNA sequencing data do not conceptually differ from the analysis of biopsy-based DNA sequencing, there are several details that require specialized bioinformatics tools. First, the amount of ctDNA in plasma or other body fluids can be very low (Heitzer *et al.*, 2019), especially following treatment, which makes variant detection extremely challenging. This caveat has motivated several approaches for error-suppression when sequencing cfDNA, which can enhance the sensitivity of variant detection (Newman *et al.*, 2016; Deng *et al.*, 2018). Another approach to go beyond the variant detection limit is to monitor previously identified mutations from the tumor rather than de novo detection. This personalized testing approach can help in determining the ctDNA detection status in a given liquid biopsy, which can be clinically relevant (Tie *et al.*, 2019). Second, cfDNA released into circulation is naturally fragmented by biological processes (Van Der Pol and Mouliere, 2019). Therefore, studying the fragmentation patterns of cfDNA is biologically relevant. The ctDNAtools package offers solutions for these two aspects of ctDNA analyses.

## 2 Methods and Implementation

The ctDNAtools package is built mainly on the *Rsamtools* and *GenomicAlignments* packages to parse aligned sequencing data in bam format. The inputs for all the functions in ctDNAtools are one or more of a bam file, mutations data frame, genomic targets data frame, and a reference genome. Figure 1 shows flowcharts of the two modules implemented in ctDNAtools.

**Figure 1:**
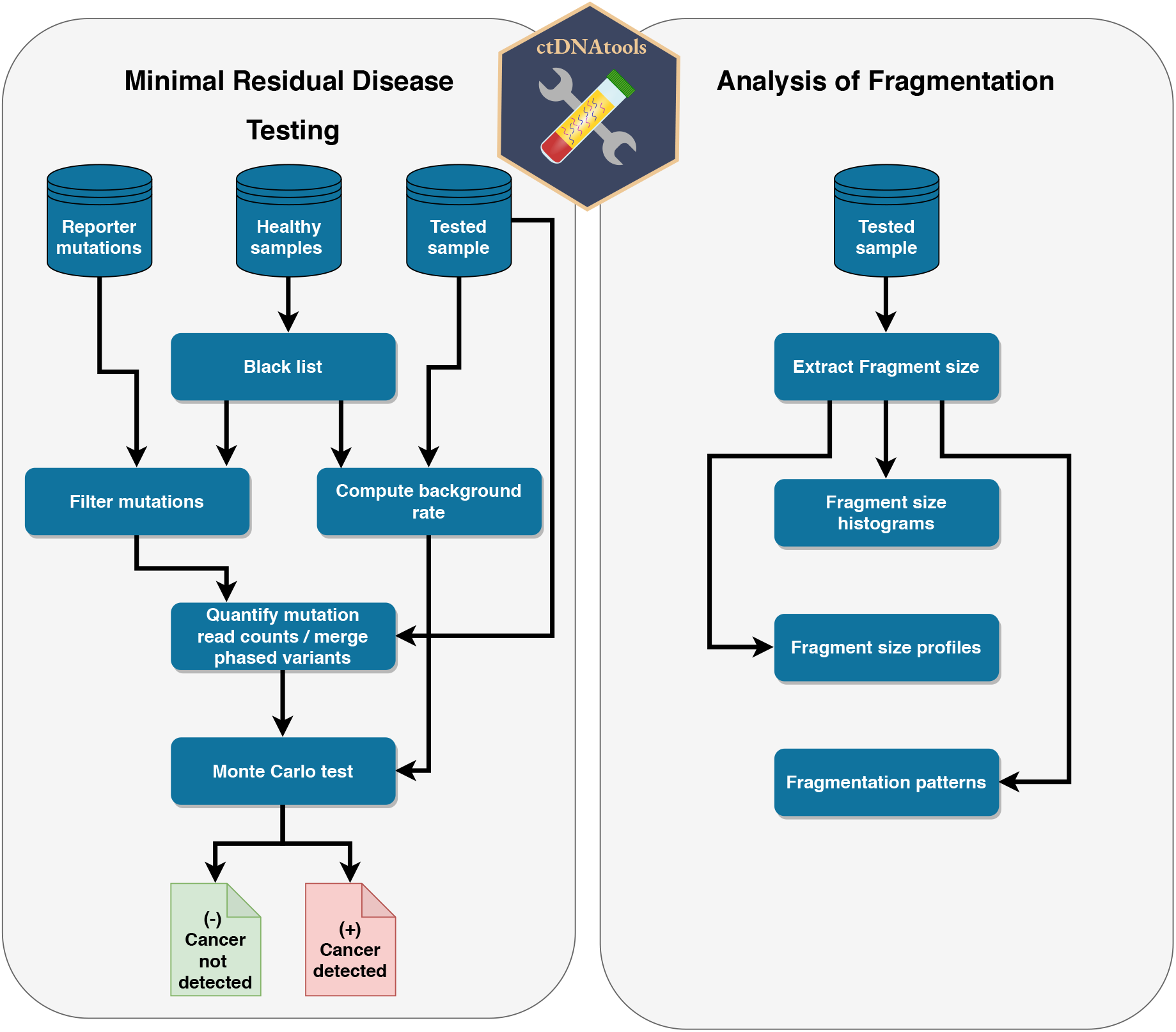
Flowcharts of the two modules in the ctDNAtools package

### 2.1 Minimal residual disease (MRD) testing

MRD is defined as the existence of cancer cells persisting in the patient after treatment at undetectable levels by imaging techniques (Pantel and Alix-Panabières, 2019). MRD testing in the ctDNAtools package is based on tracking pre-detected reporter mutations (single nucleotide variants) in a follow-up sample. The reporter mutations can be for example detected in tissue and/or liquid biopsies upon diagnosis. MRD positive samples are expected to have traces of the reporter mutations at a level that cannot be expected by chance given the mismatch rate in the tested sample. The MRD framework in ctDNAtools first counts the reads supporting reference and variant allele of the reporter mutations. Then, sample-specific mismatch rate is measured in the target regions of the sequencing panel. Finally, a Monte Carlo based sampling test is conducted to derive an empirical p-value corresponding to the sample ctDNA positivity under the background rate. There are also two optional modules that can be further incorporated including the usage of a black list of genomic loci built using a panel of control samples, and utilization of phased variants for further error suppression.

#### 2.1.1 The Monte Carlo test

Given *N* reporter mutations each with depth *D*_*i*_, in each Monte Carlo round, we randomly sample variant allele reads *X*_*i*_ under the background rate *p* using binomial distribution: *X*_*i*_ ~ *Binom*(*D*_*i*_, *p*). Then, an empirical p-value is computed based on the number of rounds where simulated data equals or exceeds observed data in two measurements: (1) the average variant allele frequency (VAF) for the *N* sampled mutations, and (2) the number of mutations with non-zero VAF. This approach was first introduced by Newman *et al.* (2016).

#### 2.1.2 Using a black list

CtDNAtools supports building a background panel from a set of controls, which can be for example plasma samples from healthy individuals. This provides a site-specific or site- and substitution-specific measurement of the mismatches in the controls. The user can then customize the threshold to define a black list of the most noisy loci in the genome. When a black list is used within the MRD module, reporter mutations within the black list will be filtered out, and the sample-specific background rate will be computed after excluding the black list. This leads to removal of likely false-positive reporter mutations and to the reduction of the mismatch rate, which typically translates to better sensitivity.

#### 2.1.3 Utilizing phased variants

Phased variants refer to closely-located somatic mutations that are exhibited in the same sequencing reads indicating a common allele. Phased variants can be found in several types of cancers forming for example double nucleotide substitutions. Particularly in lymphomas, phased variants are highly abundant as positionally clustered mutations caused by somatic hypermutation (Alkodsi *et al.*, 2019). Since these variants are always exhibited by the same reads, their traces in a follow-up sample are expected to show the same pattern (traces in the same sequencing reads). Therefore, mismatches that do not follow this pattern can be purified as confirmed artifacts, leading to advantageous reduction of the background mismatch rate. The ctDNAtools package has a functionality to merge variants in phase into a single event, while purifying the reads with confirmed artifacts. Furthermore, the background rate is adjusted with the probability of purification computed as the number of sequencing reads spanning more than one phased variant divided by the total number of reads mapping to all variants.

### 2.2 Analysis of cfDNA fragmentation

The ctDNAtools package has a function to retrieve DNA fragment lengths (*i.e.* the insert size from a bam file), while giving the user several customizable options such as mapping quality, requiring read pairs to be on different strands, or requiring reads to be properly paired. The extracted fragment lengths can be utilized in several functionalities. Fragment size histogram can be computed with ctDNAtools using fixed or custom bins, with or without normalization. Fragment size histograms differs between tumor- and normal-derived DNA, and can be used as features to machine learning algorithms for cancer detection (Mouliere *et al.*, 2018). CtDNAtools has also a function to summarize fragment sizes along defined genomic regions (fragmentation profiles), and accepts any custom summarization function the user is willing to use. These cfDNA fragmentation profiles have been shown to distinguish cancerous from healthy tissues and predict the tissue of origin (Cristiano *et al.*, 2019). Finally, ctDNAtools has a function to summarize the number of fragment ends in defined genomic regions, which reflects the underlying nucleosome occupancy (Snyder *et al.*, 2016).

### 2.3 Example data

The ctDNAtools package includes example aligned sequencing data in bam format. Two of the bam files are from sequenced plasma samples from a diffuse large B-cell lymphoma patient taken after three cycles and post-treatment. Three other bam files are from sequenced plasma samples taken from individuals with no history of malignancy or chronic illness. Sequencing data cover about 600 basepairs in chromosome 14, where 10 somatic mutations have been detected at diagnosis. These data files help the user get started with the MRD and fragmentation analyses. Supplementary Figure 1 shows the fragmentation patterns recovered from one of the samples using ctDNAtools.

## 3 Conclusion

The ctDNAtools offers a set of freely available tools to analyze ctDNA sequencing data, thereby supporting the fast development of the liquid biopsy field.

## Acknowledgements

Computational resources from CSC–IT are gratefully acknowledged.

## Supplementary figures

**Supplementary Figure 1:**
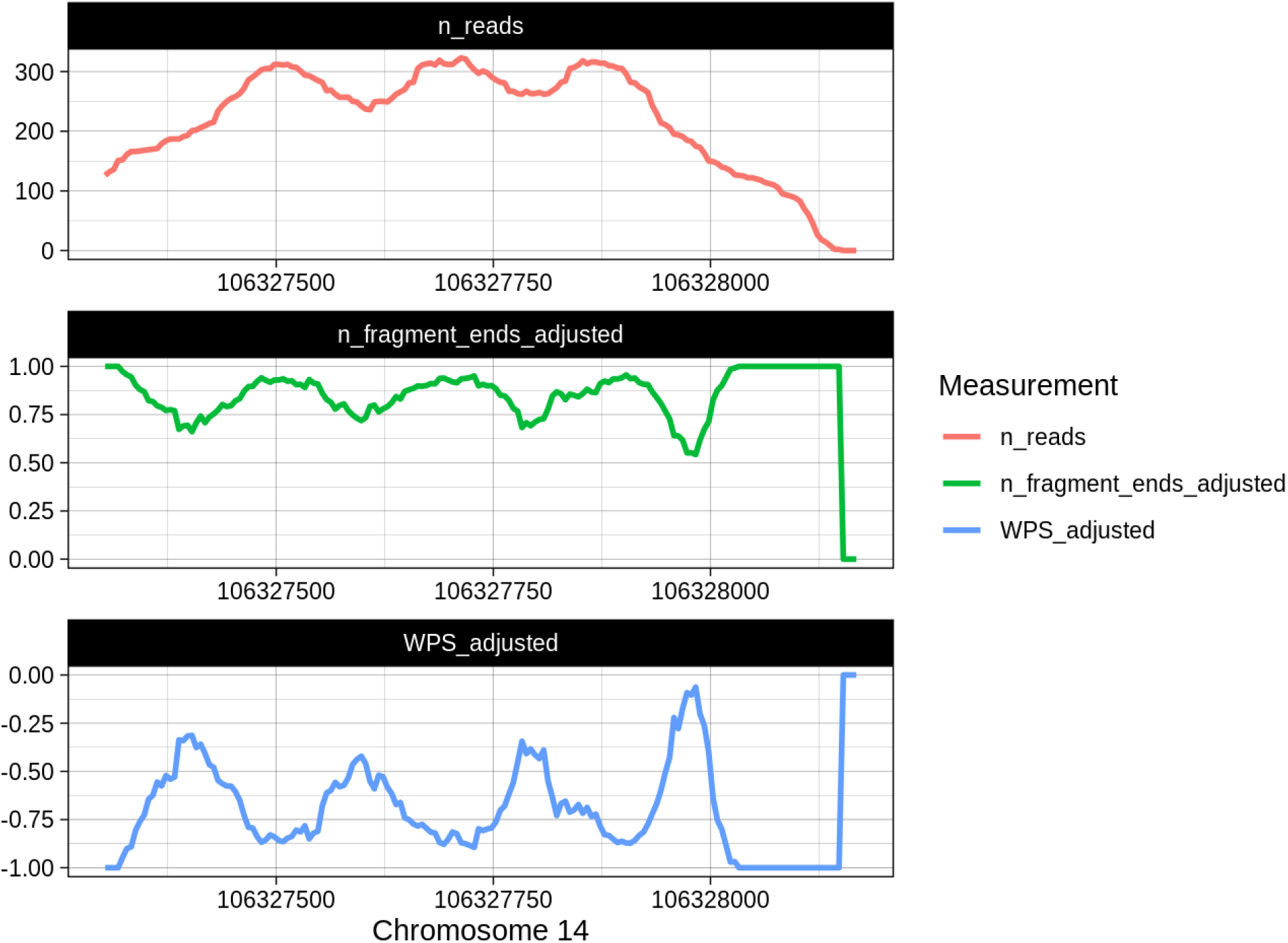
Fragmentation patterns of cfDNA extracted by ctDNAtools. The number of reads, the adjusted number of fragment ends, and the adjusted Windowed Protection Score (WPS) were computed in overlapping genomic windows of size 120 bp and step size of 5 bp. Only fragments with size 120 - 180 bp were used in this computation similar to a previous study Snyder et al. (2016)

